# Temporal Interference Stimulation of the Motor Cortex Produces Frequency-Dependent Analgesia

**DOI:** 10.64898/2026.04.15.718797

**Authors:** Amin Dehghani, David M. Gantz, Ethan K. Murphy, Ryan J. Halter, Tor D. Wager

**Author notes:** Please address correspondence to: Tor D. Wager Diana L. Taylor Distinguished Professor Presidential Cluster in Neuroscience and Department of Psychological and Brain Sciences, HB 6207 Dartmouth College Hanover, NH 03755.

## Abstract

**Background:** Transcranial temporal interference stimulation (tTIS) is an emerging noninvasive neuromodulation approach that enables focal, frequency-specific modulation of deep brain regions, offering a novel method for investigating therapeutic mechanisms underlying brain and mental health disorders. Pain is a key target because it is a feature of multiple disorders and is increasingly understood to depend on brain circuits. Here, we tested the effects of tTIS on bilateral evoked pain, capitalizing on converging evidence from human and animal studies indicating that the primary motor cortex (M1) contains body-wide inter-effector regions and has descending projections to regions implicated in nociceptive, motivational, and autonomic processing, making it a key cortical target for pain modulation. **Methods:** We conducted a pre-registered, triple-blind, randomized crossover study (N = 32, 160 study sessions), investigating frequency-dependent effects of tTIS applied to the left M1 on experimentally evoked thermal pain in healthy adults. We tested four stimulation frequencies (10 Hz, 20 Hz, 70 Hz, and sham) on separate days (>10,000 pain trials total). Noxious heat was applied to both the right and left forearms using individually calibrated temperatures both pre- and post-stimulation. **Results:** Active tTIS produced significant analgesia at all stimulation frequencies (10 Hz, 20 Hz, and 70 Hz) relative to sham (Cohen’s d = 0.46-0.82, all p < 0.05). 10 Hz produced the greatest reduction (d = 0.82), and both 10 Hz and 20 Hz produced more analgesia than 70 Hz (d = 0.44 and 0.38, respectively; p < 0.05). Stimulation-related sensations were equivalent across frequencies, and participants were blind to condition. Pain reductions remained stable over a ∼40-min post-stimulation period and were bilateral, consistent with stimulation of body-wide inter-effector regions. **Conclusions:** These results provide the first evidence that tTIS can reliably reduce experimental pain perception in humans in a frequency-dependent manner, providing a foundation for noninvasive pain modulation with tTIS.

## Introduction

Pain is a multidimensional experience shaped by peripheral nociception, spinal processing, and distributed cortical–subcortical networks (1,2). Along with reward, it is also a fundamental affective process that induces neuroplasticity, drives learning, and shapes behavior (3,4). Clinically, the brain is increasingly understood to play a central role in chronic pain (5–8). Hypersensitivity to evoked pain is a feature of many chronic pain conditions and is a key indicator of central sensitization, which is thought to contribute to the generation and maintenance of pain through multiple mechanisms (6). Beyond these disorders, pain is a central motivational function that is altered across conditions including substance use (9), depression (10), and anxiety disorders (11), and is a critical motivator of decision-making (12,13) in everyday life.

Despite advances in pharmacological and behavioral approaches, many individuals continue to experience inadequate pain relief or adverse effects from available treatments (14,15), motivating the need to better understand the brain circuitry underlying pain control and how it may be harnessed in new treatments. Neurostimulation-based approaches are emerging as promising interventions for chronic neuropathic and complex pain (16,17). Together, these considerations have motivated the search for noninvasive neuromodulation strategies that can target pain-related circuits safely, reproducibly, and with minimal side effects.

Surprisingly, one of the most promising emerging targets for pain control is the primary motor cortex (M1) (18–20). Recent retroviral tracing studies show that M1 projects directly to cortical, subcortical, and brainstem systems involved in sensory, affective, and autonomic regulation, including key pain-modulatory circuits (Figure 1a) (21). Chemogenetic activation of M1 produces layer-specific suppression of pain and avoidance behavior in mice via descending projections to the periaqueductal gray (PAG), mediodorsal thalamus, and zona incerta, without prominent motor effects (Figure 1b–c) (22). Complementary multisynaptic tracing studies in primates reveal convergence of nociceptive, motor, and autonomic control systems within M1 (23,24). In parallel, recent precision functional mapping fMRI in humans has identified distinct “inter-effector” regions embedded within M1 and interposed between traditional somatotopically mapped motor regions (25). The inter-effector regions respond in a body-wide manner and have been proposed as the cortical core of a distributed “somato-cognitive action network” (SCAN) (25). They connect preferentially to the anterior cingulate and medial thalamus, regions involved in pain, motivated decision-making, and action policy (26,27), and to subcortical and cerebellar territories implicated in arousal and autonomic regulation (Figure 1d (25)). These regions have recently been proposed as components of an “action-mode network” (AMN) (28) that regulates both behavior and autonomic function. Together, these anatomical and functional relationships motivate a circuit-level model in which M1 stimulation primarily engages SCAN inter-effector nodes, with downstream influence on motivated behavior and associated modulation of body-wide nociceptive signals in service of contextually appropriate behavior (Figure 1).

**Figure 1.**
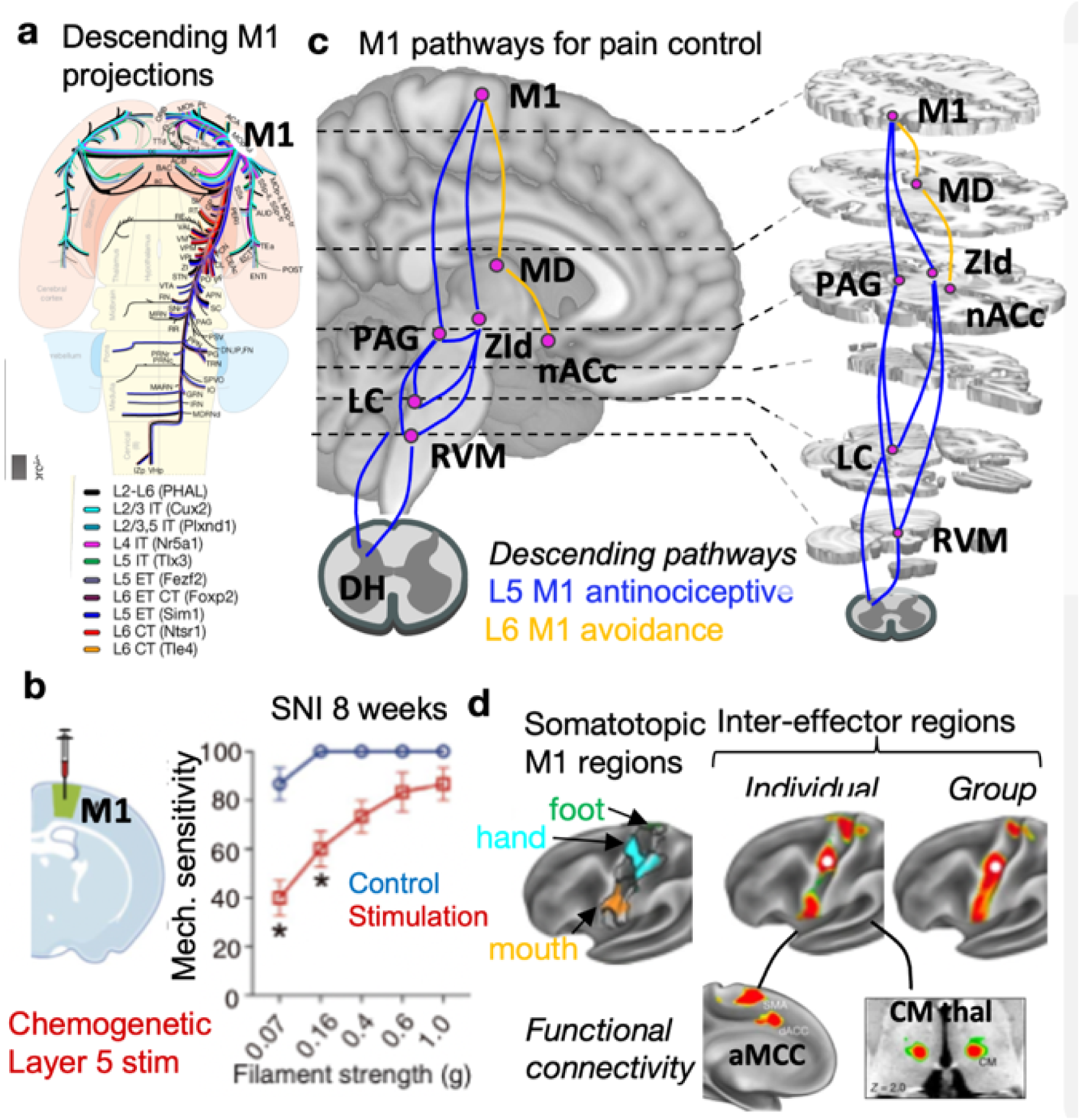
Primary motor (M1) pathways for pain control. (a) Mouse M1 contains ∼2 dozen differentiated projections targeting diverse cortical and subcortical areas, including those critical for descending pain control like the periaqueductal gray (PAG), zona incerta (ZI), and intralaminar thalamus (including centromedian CM); (21). (b) In Gan et al. (2022), chemogenetic stimulation of layer 5 M1 excitatory projections produced mechanical and cold analgesia in a spared nerve injury (SNI) neuropathic pain model. (c) Descending M1 projections to PAG and dorsal ZI mediated reduced pain hypersensitivity (blue), and a separate layer 6 M1 pathway to medial thalamus (MD) and nucleus accumbens core (nACc) reduced pain-avoidance behavior (orange). There were no motor effects of stimulating these pathways. LC = locus coeruleus, RVM = rostral ventral medulla. (d) (25) used precision fMRI mapping to identify body-wide inter-effector regions interposed between somatotopic M1 regions. These regions connect with anterior cingulate (aMCC), CM, and other cingulo-opercular network regions, and are identifiable based on functional and structural projections.

Here, we test the hypothesis that transcranial stimulation of human M1 over the approximate location of the middle inter-effector region (Figure 1d) elicits analgesia across body sites. If inter-effector regions with body-wide connectivity are important for pain control, unilateral stimulation should elicit bilateral reductions in pain. Previous studies have provided preliminary evidence that human M1 stimulation using repetitive transcranial magnetic stimulation (rTMS) (29,30), transcranial direct current stimulation (tDCS) (31–33), and transcranial alternating current stimulation (tACS) (34,35) can be analgesic in both clinical pain conditions and experimentally evoked pain (36–38). However, few studies have tested effects on multiple body sites (31,39–43), as needed to test the inter-effector hypothesis. In addition, prior reviews and meta-analyses of tDCS and tACS have provided mixed results, with some failures to find effects (18,38,44–46), and methodological limitations with these techniques may be a barrier to finding consistent effects. Thus, the role of inter-effector regions in human pain control remains unclear.

Our preregistered randomized crossover trial (47) stimulated left M1 using Transcranial Temporal Interference stimulation (tTIS), a recently developed noninvasive method designed to address limitations of conventional transcranial electrical stimulation techniques (48–50). By delivering two high-frequency (≥1 kHz) currents that interact to generate a low-frequency amplitude-modulated envelope, tTIS enables frequency-specific neural modulation of cortical and deep brain structures. Computational modeling (51,52), non-human animal (48,53), and human studies (54,55) indicate that tTIS can achieve greater effective depth than other transcranial electromagnetic approaches and modulate regional activity in a frequency-dependent manner (56,57). It also stimulates a larger envelope than TMS, which is highly sensitive to the precise coil position, and is thus more likely to robustly modulate inter-effector regions across participants. Participants received M1-targeted tTIS at envelope frequencies of 10, 20, and 70 Hz, as well as sham stimulation. Effects of tTIS were assessed using individually calibrated noxious heat stimuli applied to both the right and left forearm, maximizing sensitivity to stimulation-related changes while reducing between-participant variability. If M1 tTIS elicits bilateral analgesia, this would provide causal evidence that motor pathways influence human pain via non-somatotopic (i.e., inter-effector) pathways.

## Results

Participants (n = 32, M_age_ = 24.7 ± 4.6, range: 18–34 years, 16 biological female and 16 biological male) experienced calibration followed by 4 sessions (2–5) with the 4 treatments (sham, 10, 20, and 70 HZ tTIS) in counterbalanced order on different days, with at least 4 days between sessions (160 total sessions; see Methods; preregistration on Open Science Framework (47)).

## Calibration

Thermal pain and tolerance thresholds were 46.5 ± 0.9°C and 48.3 ± 0.8°C (mean ± standard deviation; see Figure 2(d)). There were no significant sex differences in pain threshold or tolerance (p > 0.2).

**Figure 2.**
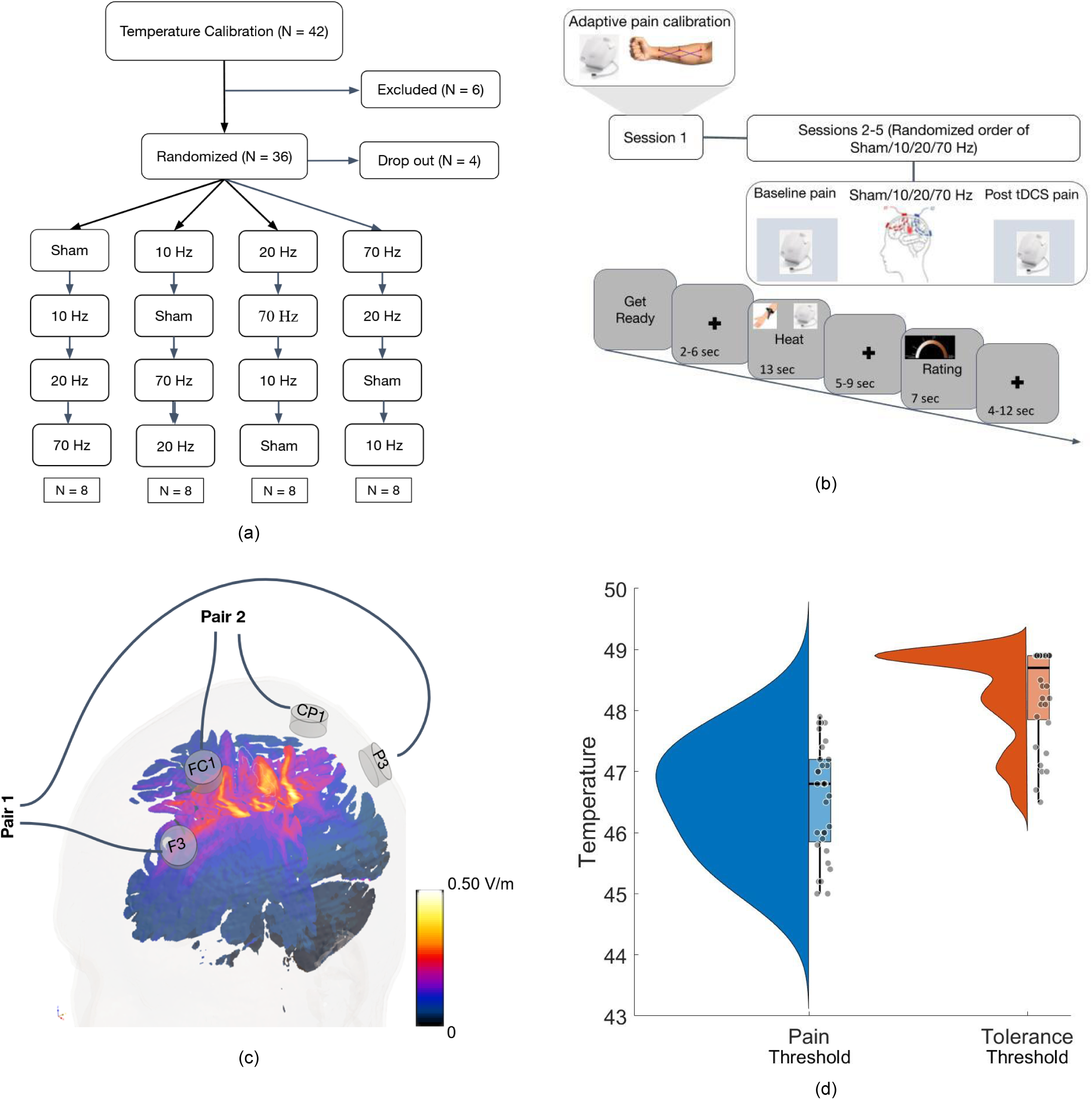
Experimental design, stimulation montage, and pain calibration. (A–B) Participant flow and randomized within-subject crossover design. Following temperature calibration, participants completed four stimulation sessions (sham, 10, 20, and 70 Hz tTIS) in randomized order on separate days. Each trial lasted 40 s and began with a warning cue (“Get Ready”; 1 s), followed by a 2–6 s delay, noxious heat stimulation (13 s total, including 1.5 s ramp-up/down and a 10 s plateau), a 5–9 s delay, a pain rating period (7 s, using the right hand), and a final 4–12 s inter-trial interval. Pain ratings were provided using a semi-circular general Labeled Magnitude Scale (gLMS). (c) Electric field distribution produced by an optimized tTIS montage targeting the left primary motor cortex (M1), using F3–P3 and CP1–FP1 electrode pairs, as estimated by a computational model. The model illustrates electrode placement and the resulting peak field strength. (d) Distributions of individually calibrated temperatures used to elicit pain and tolerance across participants (mean ± SD: 46.5 ± 0.9 °C for pain and 48.3 ± 0.8 °C for tolerance).

## Sensations caused by tTIS

In sessions 2–5, following each tTIS session, participants rated the sensations evoked by stimulation. Ratings included itching, tingling, metallic/iron taste, pleasantness or unpleasantness, warmth/heat, and pain (see Figure 3). A one-way repeated-measures ANOVA comparing sham, 10 Hz, 20 Hz, and 70 Hz stimulation sessions revealed no statistically significant differences in reported tTIS sensations across stimulation types (all p-values > 0.05), indicating no evidence that the type of stimulation systematically influenced sensory experiences. Analyses of blinding indicated that participants were blind to stimulation conditions (see “Assessments of blinding” below).

**Figure 3.**
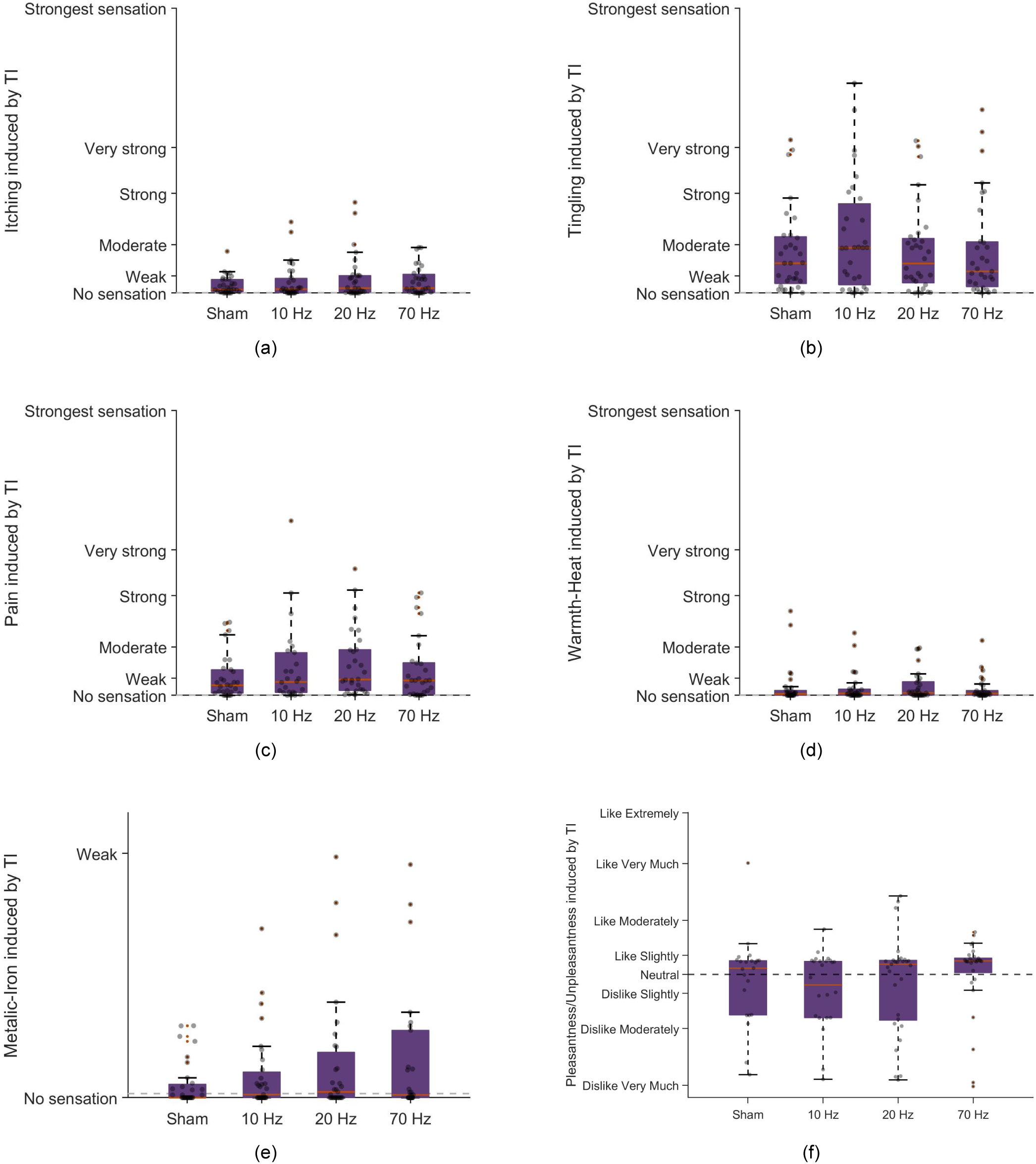
Sensory experiences during tTIS. Box-and-whisker plots of Temporal Interference-induced sensations of (a) itch, (b) tingling, (c) pain, (d) warmth, (e) metallic-iron taste, (f) pleasant vs. unpleasant sensations. Tingling was the most strongly endorsed sensation, which was slightly unpleasant. Importantly, none of the sensations differed significantly across sham, 10, 20, and 70 Hz conditions (all p > 0.05).

To characterize the underlying structure of participants’ sensory experiences during stimulation, we performed an exploratory factor analysis on the six rated dimensions. Two latent factors emerged. The first factor, marked by strong positive loadings for itching, warmth, metallic sensation, and tingling, represents a somatosensory–physical sensation dimension, capturing general cutaneous and bodily sensations commonly associated with tTIS. The second factor, defined by a positive loading for pain and a negative loading for pleasantness, reflects an affective–aversive dimension, with higher values reflecting aversiveness. Factor scores for both dimensions did not differ significantly across sham and active stimulation conditions (all p > 0.05; see Methods). These two factors were subsequently used in secondary analyses to assess whether specific sensory experiences moderated the effects of tTIS on pain outcomes (see “Sensitivity analysis” below).

## Effects of stimulus intensity and time on pain

Effects of tTIS Condition (10 Hz, 20 Hz, 70 Hz, Sham), Body Side (ipsi- vs. contralateral, fully crossed, within condition), stimulus intensity (fully crossed, within condition), and Run within testing session and Trial within run (fully crossed, within condition) were tested in a single linear mixed effects model with post-treatment pain as outcome and mean pre-treatment pain per session as a covariate (58,59), in keeping with our preregistration, prior published work (60,61), and published methodological recommendations (58–61). Treatment order was included as a between-person covariate. Stimulation effects were tested with planned contrasts (Active vs. Sham, 10 Hz vs. the average of 20 and 70 Hz, and 20 vs. 70 Hz) Generalizability across Body Side was tested by including the tTIS Condition x Body Side interaction. Generalizability across stimulus intensities and durability over time were tested with two-way interactions tTIS Condition x Stimulus Intensity and tTIS Condition x Run, respectively. Effects of biological sex and age were tested in an extended model (see Sensitivity Analysis below). Full fixed-effect coefficients, F-statistics, error degrees of freedom, and P values for the main model (see Equation 1) are reported in Table 1.

**Table 1.**
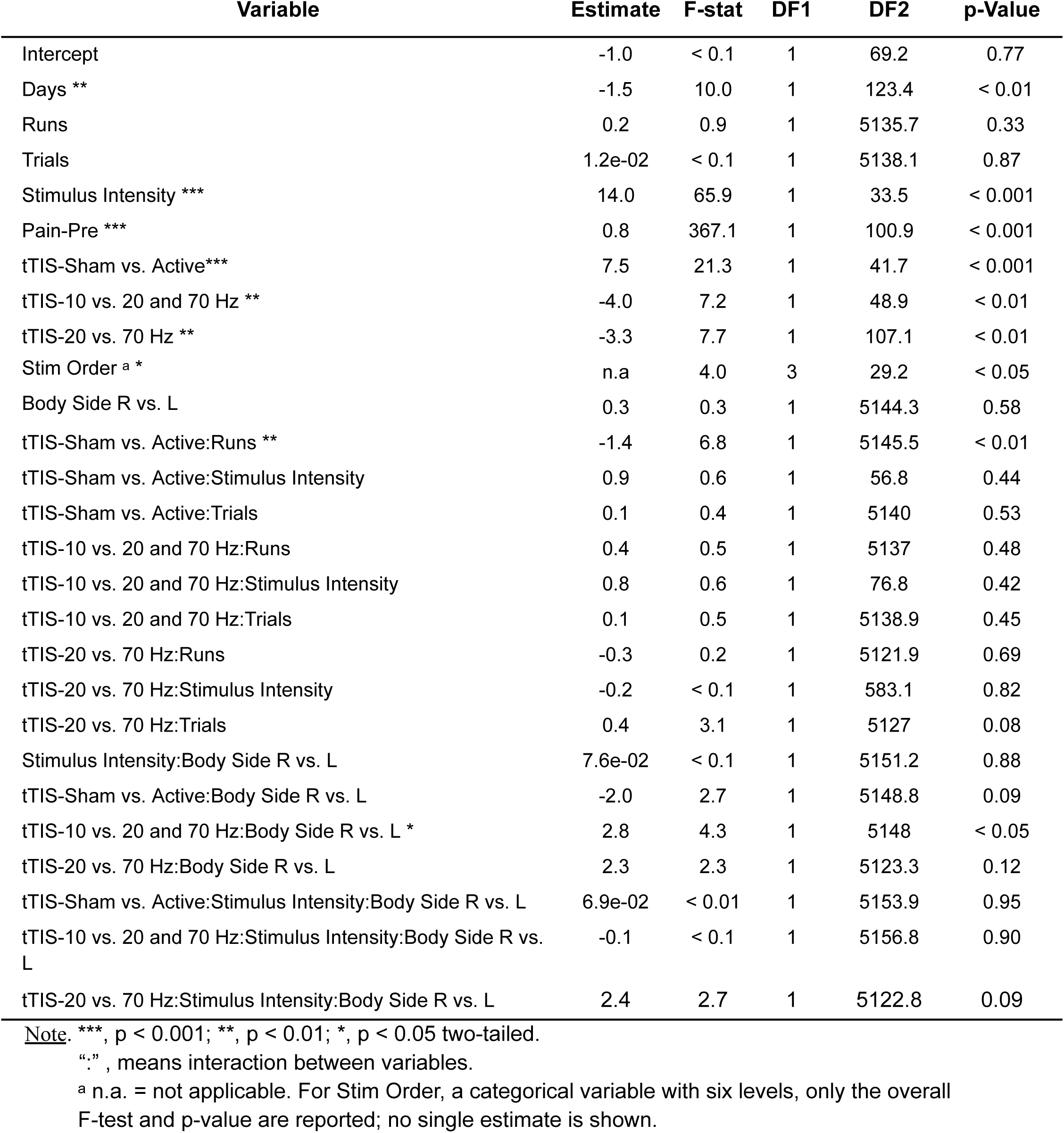
Mixed-effects model results for pain perception after tTIS.

| Variable | Estimate | F-stat | DF1 | DF2 | p-Value |
| --- | --- | --- | --- | --- | --- |
| Intercept | -1.0 | < 0.1 | 1 | 69.2 | 0.77 |
| Days ** | -1.5 | 10.0 | 1 | 123.4 | < 0.01 |
| Runs | 0.2 | 0.9 | 1 | 5135.7 | 0.33 |
| Trials | 1.2e-02 | < 0.1 | 1 | 5138.1 | 0.87 |
| Stimulus Intensity *** | 14.0 | 65.9 | 1 | 33.5 | < 0.001 |
| Pain-Pre *** | 0.8 | 367.1 | 1 | 100.9 | < 0.001 |
| tTIS-Sham vs. Active*** | 7.5 | 21.3 | 1 | 41.7 | < 0.001 |
| tTIS-10 vs. 20 and 70 Hz ** | -4.0 | 7.2 | 1 | 48.9 | < 0.01 |
| tTIS-20 vs. 70 Hz ** | -3.3 | 7.7 | 1 | 107.1 | < 0.01 |
| Stim Order <sup>a</sup> * | n.a | 4.0 | 3 | 29.2 | < 0.05 |
| Body Side R vs. L | 0.3 | 0.3 | 1 | 5144.3 | 0.58 |
| tTIS-Sham vs. Active:Runs ** | -1.4 | 6.8 | 1 | 5145.5 | < 0.01 |
| tTIS-Sham vs. Active:Stimulus Intensity | 0.9 | 0.6 | 1 | 56.8 | 0.44 |
| tTIS-Sham vs. Active:Trials | 0.1 | 0.4 | 1 | 5140 | 0.53 |
| tTIS-10 vs. 20 and 70 Hz:Runs | 0.4 | 0.5 | 1 | 5137 | 0.48 |
| tTIS-10 vs. 20 and 70 Hz:Stimulus Intensity | 0.8 | 0.6 | 1 | 76.8 | 0.42 |
| tTIS-10 vs. 20 and 70 Hz:Trials | 0.1 | 0.5 | 1 | 5138.9 | 0.45 |
| tTIS-20 vs. 70 Hz:Runs | -0.3 | 0.2 | 1 | 5121.9 | 0.69 |
| tTIS-20 vs. 70 Hz:Stimulus Intensity | -0.2 | < 0.1 | 1 | 583.1 | 0.82 |
| tTIS-20 vs. 70 Hz:Trials | 0.4 | 3.1 | 1 | 5127 | 0.08 |
| Stimulus Intensity:Body Side R vs. L | 7.6e-02 | < 0.1 | 1 | 5151.2 | 0.88 |
| tTIS-Sham vs. Active:Body Side R vs. L | -2.0 | 2.7 | 1 | 5148.8 | 0.09 |
| tTIS-10 vs. 20 and 70 Hz:Body Side R vs. L * | 2.8 | 4.3 | 1 | 5148 | < 0.05 |
| tTIS-20 vs. 70 Hz:Body Side R vs. L | 2.3 | 2.3 | 1 | 5123.3 | 0.12 |
| tTIS-Sham vs. Active:Stimulus Intensity:Body Side R vs. L | 6.9e-02 | < 0.01 | 1 | 5153.9 | 0.95 |
| tTIS-10 vs. 20 and 70 Hz:Stimulus Intensity:Body Side R vs. L | -0.1 | < 0.1 | 1 | 5156.8 | 0.90 |
| tTIS-20 vs. 70 Hz:Stimulus Intensity:Body Side R vs. L | 2.4 | 2.7 | 1 | 5122.8 | 0.09 |

Note. \*\*\*, $p < 0.001$ ; \*\*, $p < 0.01$ ; \*, $p < 0.05$ two-tailed.
“:”, means interaction between variables.
<sup>a</sup> n.a. = not applicable. For Stim Order, a categorical variable with six levels, only the overall F-test and p-value are reported; no single estimate is shown.

Several methodological variables produced expected effects on pain. All were orthogonal to treatment condition by design and additionally controlled for in the analysis of stimulation effects described below. As expected, pain was significantly higher at higher Stimulus Intensity (F(1,33.5) = 65.9, p < 0.001), and post-treatment pain was strongly predicted by pre-treatment pain (Pain-Pre: F(1,100.9) = 367.1, p < 0.001). These provide positive controls for pain data quality and reliability.

Pain decreased significantly across the four treatment Days (F(1,123.4) = 10.0, p < 0.01), indicating overall habituation, but did not vary across Runs within day (F(1,5135.7) = 0.9, p = 0.33) or Trials within run (F(1,5138.1) < 0.1, p = 0.87). The main effect of treatment order was significant (F(3,29.2) = 4.0, p < 0.05), indicating modest differences in overall pain sensitivity across the four counterbalanced stimulation orders. These were orthogonal to treatment conditions because we counterbalanced the assignment of treatments to days across participants (Figure 2; see Methods).

## tTIS effects on pain

### Effects of active vs. sham stimulation

As shown in Figure 4 (a) and Table 1, post-treatment pain was significantly reduced after active tTIS vs. Sham (tTIS-Sham vs. Active, averaging across stimulation frequencies); F(1,41.7) = 21.3, p < 0.001). Planned contrasts revealed that each individual tTIS frequency produced significant analgesia compared with Sham (all P < 0.05) with moderate to large effect sizes (10 Hz vs. Sham: *d* = -0.82 (95% CI [-1.22, -0.42]), 20 Hz vs. Sham: d = -0.60 (CI [-0.97, -0.22]), 70 Hz vs. Sham: d = -0.46 (CI [-0.83, -0.10]). In addition, pain decreased significantly for 10 Hz vs. the average of 20 and 70 Hz (F(1,48.9) = 7.2, p < 0.01) and 20 vs. 70 Hz (F(1,107.1) = 7.7, p < 0.01). Thus, 10 Hz stimulation was most effective, with higher frequencies producing relatively weaker analgesia.

**Figure 4.**
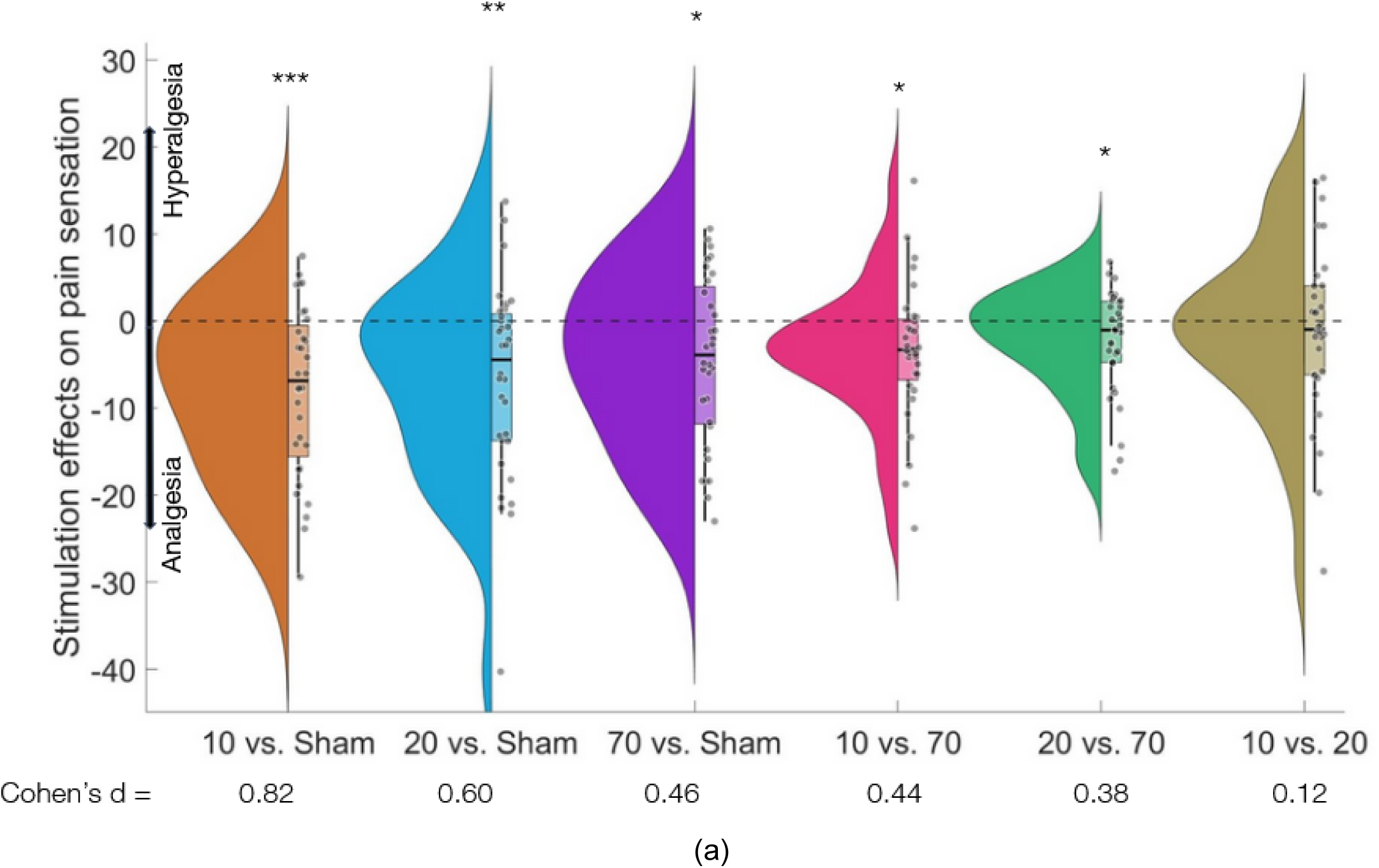
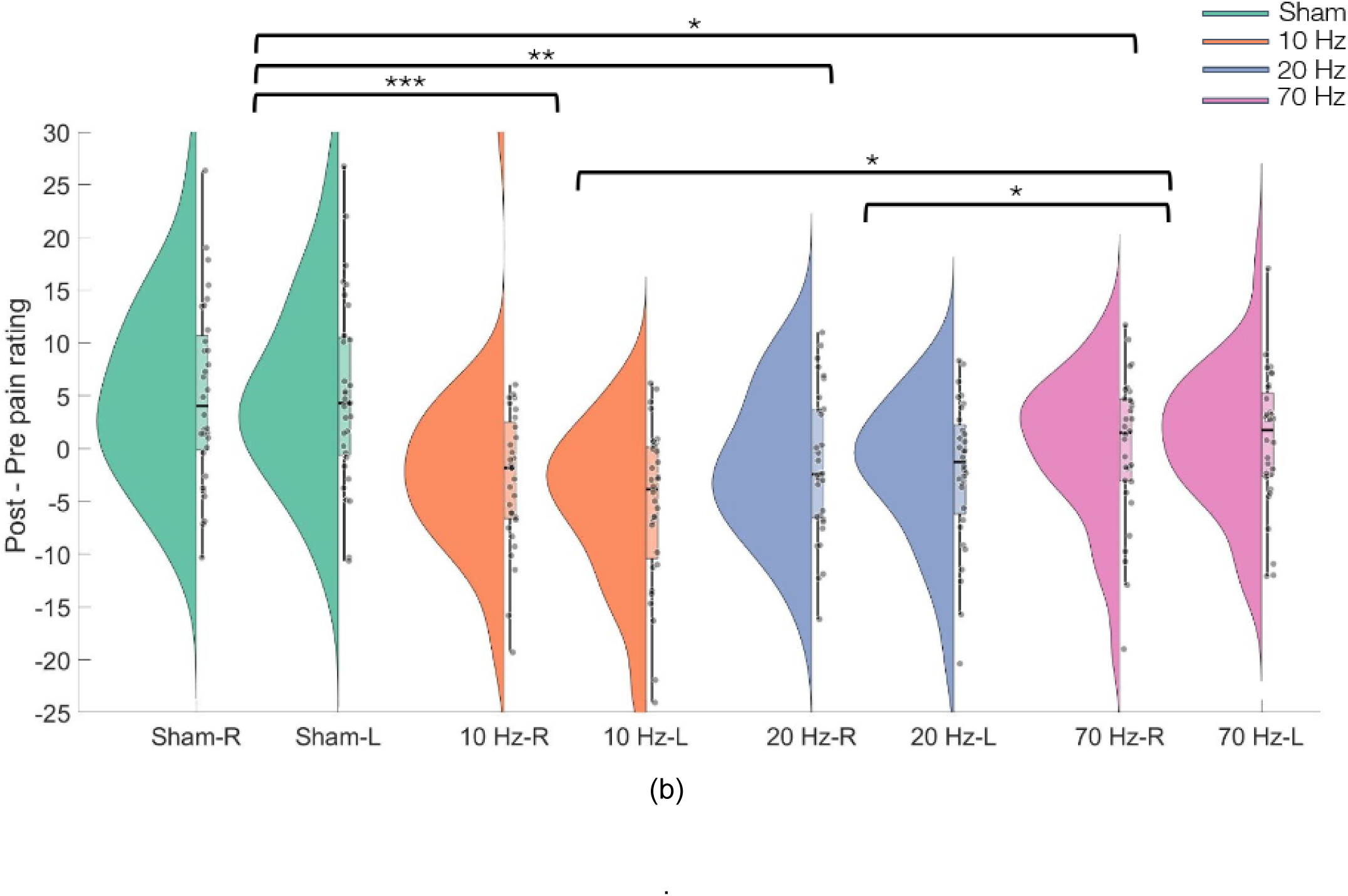
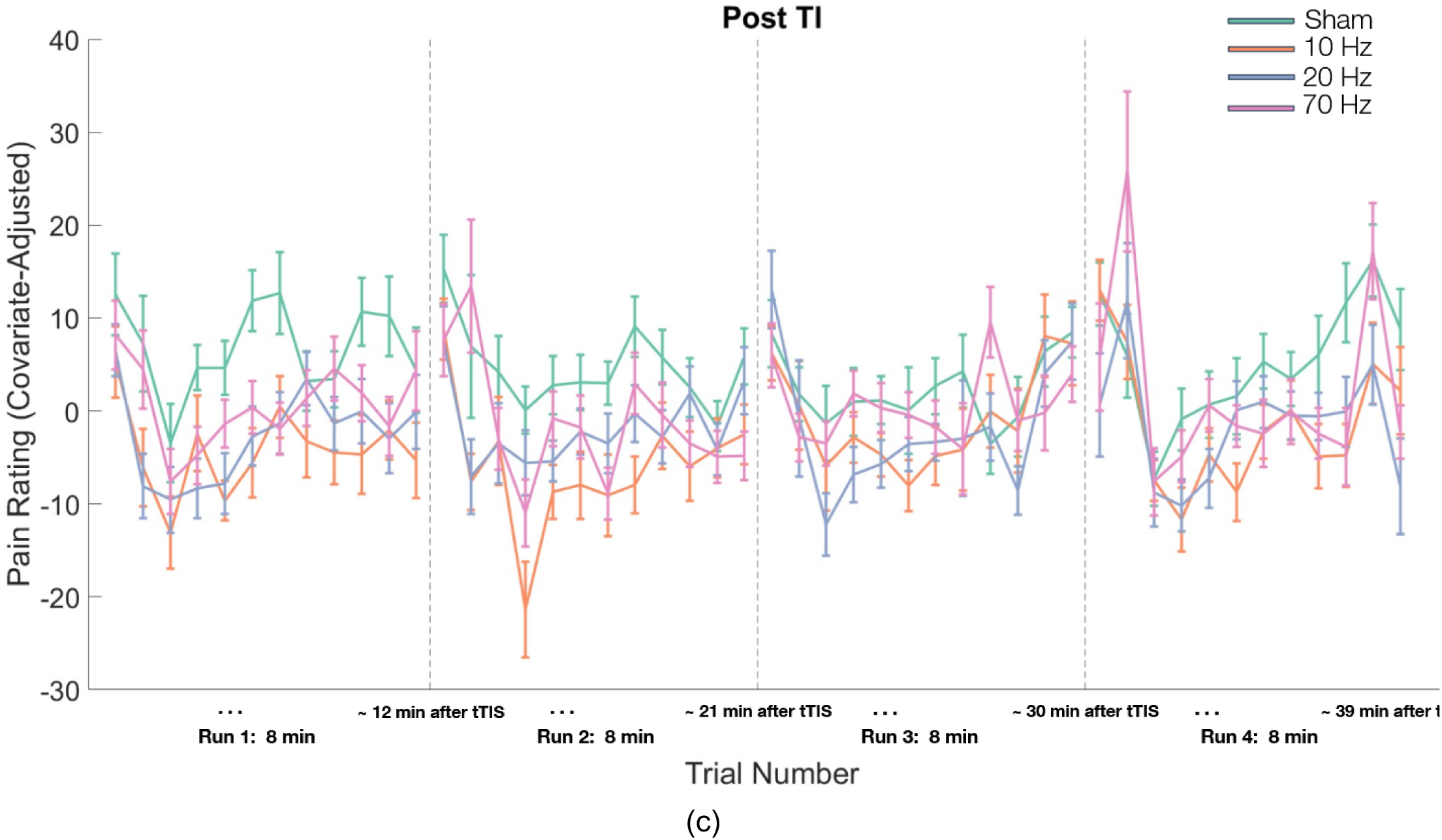
Main findings for pain sensation. (A) The effects of 10, 20, 70 Hz vs. sham, 10 and 20 vs. 70 Hz, and 20 vs. 70 Hz on pain sensation. (B) Treatment effects on left and right hand (Post-treatment—Pretreatment pain rating change) for each of the 4 stimulation types. (C) Model-corrected prestimulation pain ratings across 4 runs and 12 trials within each run for 10, 20, 70 Hz, and sham stimulations. Corrections removed variance from nuisance covariates (eg, Pain-Pre, Days, Stimulus Intensity, Stim Order, Age, Body Side, Biological Sex) using a simplified mixed-effects model. This isolates stimulation-related effects on pain trajectories. Boxplots indicate the median, 25th, and 75th percentiles, and individual data points. Dots reflect effect magnitude estimates of individual participants. All values were adjusted to remove participant-level intercepts. \*\*\**P* < 0.001; \*\**P* < 0.01; \**P* < 0.05 two-tailed.

On the generalized labeled magnitude scale (gLMS), low-intensity stimuli in the sham condition were rated at 46.5 before stimulation and 44.0 after stimulation, corresponding to sensations between moderate and strong (see *Adaptive Thermal Calibration Procedure (Session 1)* in Methods), with a 2.4% reduction. At 10 Hz, low-intensity ratings decreased from 48.8 to 38.9, remaining within the moderate–strong range but shifting toward moderate (20.7% reduction). At 20 Hz, ratings decreased from 47.2 to 38.5, also remaining within the moderate–strong range (18.9% reduction). At 70 Hz, ratings decreased from 48.5 to 42.4, remaining within the moderate–strong range (10.9% reduction).

For high-intensity stimuli, sham ratings decreased from 76.2 to 73.3, both corresponding to gLMS sensations between strong and very strong, with a 3.5% reduction. At 10 Hz, ratings decreased from 78.9 to 67.5, remaining within the strong–very strong range but shifting toward strong (14.2% reduction). At 20 Hz, ratings decreased from 76.7 to 66.5, also remaining within the strong–very strong range but shifting toward strong (13.5% reduction). At 70 Hz, ratings decreased from 76.6 to 68.4, remaining within the strong–very strong range (10.2% reduction).

### Laterality of stimulation effects

Pain ratings did not differ between the right (contralateral) and left (ipsilateral) forearms overall (post-pain across conditions, F(1,5144.3) = 0.3, p = 0.58), and there was no significant Active vs. Sham x Body Side interaction (F(1,5148.8) = 2.7, p = 0.09). The numerical direction favored greater ipsilateral than contralateral analgesia with active stimulation. In addition, the interaction between stimulation frequency (10 Hz vs. 20 and 70 Hz) and Body Side was significant (F(1,5148) = 4.3, p < 0.05, with higher analgesia on the left (ipsilateral) forearm vs. right (contralateral) forearm. This finding indicates that 10 Hz produced relatively greater analgesia on the ipsilateral forearm vs. contralateral forearm. Thus, frequency effects were not bilaterally symmetrical. However, the effect contrasts with typical stimulation effects on motor function, which are mainly contralateral (20,62). Together, these findings are consistent with bilateral analgesia, and by extension the inter-effector hypothesis (see Figure 4b). Pain rating changes for high-temperature and low-temperature conditions across all stimulation types are presented in Figure S1 (see Supplementary Materials).

### Durability of stimulation effects

Active vs. sham tTIS effects generally persisted across the testing session (Figure 4). However, a significant Sham vs. Active x Run interaction (F(1,5145.5) = 6.8, p < 0.01) consistent with a modest reduction of the effect across the test session. However, interactions between the active stimulation contrasts (10 vs. 20 and 70 Hz; 20 vs. 70 Hz) and Runs/Trials were not significant (all p > 0.05, see Table 1), suggesting largely stable effects of tTIS frequency over time. In addition to these effects, we observed a sharp decrease in pain ratings from the first to the second trial within each run (Figure 4c), consistent with prior reports (63–65). This early reduction likely reflects rapid sensory adaptation and central habituation to repeated noxious stimulation. This effect was orthogonal to treatment effects in our experimental design, so does not confound other effects presented above, and did not impact the findings reported above (see Sensitivity Analysis below). In sum, the analgesic effects of the 20-min tTIS intervention remained relatively stable across the ∼35–40 min post-stimulation testing period.

## Sensitivity analyses to examine robustness of tTIS effects

Above, we reported effects from our pre-registered mixed effects model. To examine robustness, we conducted several types of sensitivity analysis to test whether the main findings remained significant across different modeling choices. In an extended mixed-effects model, we additionally adjusted for age and biological sex, as well as their interactions with tTIS. This model included additional random participant slopes for Day, Run, Trial, Run x tTIS condition, and Trial x tTIS condition (i.e., a random-effects specification of the form (1 + Days + (Runs + Trials + Stimulus Intensity) × tTIS | Participant), with tTIS coded using the contrasts Sham vs. Active, 10 vs. 20 and 70, and 20 vs. 70). The results showed that the effect of Sham vs. Active and the effect of 10 vs. 20 and 70 Hz remained significant (p < 0.001 and p = 0.01, respectively), but the effect of 20 vs. 70 Hz was not significant (p = 0.07). There was no significant difference in pain between males and females (Biological Sex: F(1,25.9) < 0.1, p = 0.90), but pain was significantly lower in older participants (F(1, 29.4) = 7.3, p < 0.05). There were no significant interactions between tTIS condition and either Biological Sex or age (tTIS × Sex and tTIS × Age, all p > 0.10). However, the relatively restricted age range of the sample limits the interpretation of age-related effects across the lifespan (See equation S.1).

A second model tested potential interactions with tTIS sensations, which could potentially alter expectations and thus affect pain ratings, as has been found with previous studies of medication side effects (66). The two tTIS-sensation factors (physical sensation strength and aversion) were included as predictors, along with their interactions with tTIS condition in the mixed-effects model in equation (1). Neither sensory factor showed a significant main effect on pain, nor did either significantly interact with tTIS condition (all p > 0.1). The only significant interaction was between the tTIS-10 vs. 20 vs. 70 Hz and factor 2 (p < 0.05). Importantly, the main effects of tTIS (Sham vs. Active, 10 vs. the average of 20 and 70 Hz, and 20 vs. 70 Hz) remained significant (all *p* < 0.01), indicating that the observed effects on pain were not attributable to individual differences in tTIS-related sensations across stimulation conditions.

Given the sharp drop in ratings from the first to the second trial, we tested a third variant of the primary mixed-effects model (see equation (1)) with the first trial of each run excluded from analysis. All tTIs effects remained significant, including Sham vs. Active, 10 vs. 20 and 70 Hz, and 20 vs. 70 Hz (p < 0.001, p = 0.01, and p < 0.01, respectively).

Finally, to assess potential randomization failures, a one-way repeated measures ANOVA was conducted on the average pre-tTIS pain ratings across the three stimulation sessions. The analysis revealed no significant differences in baseline pain ratings among sessions for either the low-temperature stimuli (F(2, 125) = 0.14, p = 0.87) or the high-temperature stimuli (F(2, 125) = 0.11, p = 0.89). These results suggest that randomization was adequate and there were no group differences at baseline.

## Assessments of blinding

Analyses of debriefing questions indicated that participants could not discriminate active from sham tTIS. 25 of 32 (78%) participants initially guessed that all conditions were active. Participants were then informed that exactly one session was sham and asked to identify which of the four sessions was the sham (with the option to continue to report all as active). Participants performed at chance, with 5 out of 32 (15%) guessing correctly (95% CI [7%, 31%]; chance = 20%; exact binomial test, two-sided p = 0.66).

## Reaction Time Control Analysis

The laterality of effects (bilateral or weakly ipsilateral) does not match the expected pattern of effects for stimulation of classic somatotopic motor pathways, but further tests of relationships with motor function are of interest. As an additional control analysis not specified in the preregistration, we examined reaction time during pain ratings, operationalized as time to movement initiation on the rating scale. Participants reported pain ratings using the right hand, contralateral to the stimulated left M1. If stimulation had influenced motor response processes, differences in reaction time across stimulation conditions might be expected. However, there were no significant effects of tTIS-Sham vs. Active (F(1,41.3)=1.3, p = 0.26), tTIS-10 Hz vs. the average of 20 and 70 Hz (F(1,45.6)=0.2, p = 0.62), tTIS-20 Hz vs. 70 Hz (F(1,43.3)=0.1, p = 0.70), Body Side (F(1,5153.7) < 0.1, p = 0.76), or their interaction (all p > 0.5) on reaction time. These findings indicate that stimulation did not measurably alter motor response speed. Taken together, the findings suggest that tTIS-induced analgesia is unlikely to be explained by lateralized motor effects or response facilitation associated with stimulation of the motor cortex.

## Discussion

Recent advances in the neuroscience of pain demonstrate complex control over nociception and pain behaviors by central circuitry (8,67–69). Noninvasive neuromodulation offers a promising new way to intervene at the brain circuit level (37,70). However, the effects of current noninvasive approaches are modest and variable across studies (18,38,45,46). TTIS stimulation offers great promise in its ability to deliver more current to deeper brain structures (48,53), as well as allowing for frequency-specific stimulation. This preregistered, randomized trial provides the first evidence that tTIS applied over the left M1 can reliably reduce acute thermal pain intensity ratings in humans, and that it does so in a body site-general manner, consistent with stimulation of body-wide inter-effector regions (25) and with descending M1→PAG and M1→medial thalamic projections identified in rodent models (22). Across participants, all active stimulation frequencies produced significant analgesia relative to sham with large effect sizes (up to d = 0.82), with effects strongest at 10 Hz, intermediate at 20 Hz, and smaller but reliable at 70 Hz. Analgesic effects were largely stable across the full post-stimulation period (0 - 40 min post-tTIS). The induction of bilateral analgesia and lack of effects on reaction times are both consistent with body-wide inter-effector regions as key targets rather than lateralized motor representations. These effects were not explained by differences in perceived sensations or unblinding. The triple-blind (participant, experimenter, and analyst) crossover experimental design eliminated confounding of effects with baseline pain, temporal habituation or sensitization, participant expectancy, and experimenter expectancy. In addition, participants were unable to distinguish active from sham stimulation, indicating adequate blinding. Sensitivity analyses confirmed the robustness of tTIS effects to various analytic choices.

The frequency dependent pattern of effects (10 Hz > 20 Hz > 70 Hz) provides clues on potential mechanisms. Pain arises from distributed and interacting neural systems that encode sensory processing, affective responses, expectations, and prediction errors (3,71,72), and oscillations across frequency bands are thought to coordinate communication within and between these systems (4,73,74). Although frequency bands are often described as distinct, they can also interact, and alpha and beta oscillations in particular are sometimes dissociable (75) but in other contexts covary to support shared functions such as top-down control. The graded pattern observed here is therefore consistent with a frequency-dependent engagement of overlapping neural mechanisms rather than strictly frequency-specific effects.

The strong analgesic effect at 10 Hz, generally considered in the “high alpha” range, is consistent with the role of alpha rhythms in inhibitory control and sensory suppression (76). Increased alpha activity is associated with reduced pain (77–79), and alpha stimulation enhances inhibitory tone in sensorimotor and prefrontal networks (80,81). Individuals with higher peak alpha frequency show lower experimental pain (82,83) and reduced facilitation of spinal (i.e., nociceptive flexion reflex) and supraspinal (i.e., N2 potential) nociception by threatening images (84) (but cf. (85,86)), whereas prolonged pain reduces peak alpha frequency and high alpha (10–12 Hz) (87). Alpha activity is also linked to expectation-related modulation of pain, suggesting a role in integrating anticipatory signals with nociceptive input (88). Alpha reductions linked to pain also fit within the framework of thalamocortical dysrhythmia, in which reduced afferent input results in impaired inhibitory control (or as a shift in excitatory-inhibitory balance) (89). This pattern is thought to manifest as a shift from alpha to lower theta frequency oscillations in thalamocortical circuits coupled with surrounding gamma increases (cross-frequency coupling) (89–91), creating enhanced pain along with other sensory and motor symptoms across disorders (90). Accordingly, reductions in alpha power or frequency may diminish inhibitory control and facilitate the propagation of nociceptive activity (92–94). Our observation of stronger analgesic effects with alpha stimulation relative to other frequencies is broadly consistent with these other observations, and suggests that 10 Hz (or higher) stimulation may help establish a balance in thalamocortical circuits that favors inhibition.

TTIS analgesia at higher frequencies may capitalize on similar principles, but may also involve different mechanisms. The intermediate effect at 20 Hz suggests partial engagement of beta-related systems involved in sensorimotor integration (95) and predictive coding of pain (96), with beta reflecting top-down predictions. Beta rhythms are associated with the maintenance of sensory representations and top-down control (97), including attentional modulation of nociceptive processing, and can influence pain ratings depending on context. In line with the graded pattern observed here, beta-band effects may reflect partial recruitment of top-down control mechanisms that overlap with, but are less strongly inhibitory than, alpha-mediated processes, consistent with the idea that alpha and beta can either dissociate or jointly contribute to higher-order control depending on task demands.

The smaller effect at 70 Hz may reflect the more complex role of gamma activity in pain. On one hand, nociceptive stimuli have been shown to induce gamma oscillations in the primary somatosensory cortex whose amplitude scales with both stimulus intensity and subjective pain (98). Some studies have suggested that gamma activity may reflect the selective amplification of nociceptive signals for conscious perception (98), consistent with gamma as a source of aberrant percepts in the thalamocortical dysrhythmia model. However, prestimulus gamma has been shown to predict lower subsequent pain (79). In addition, gamma activity is also linked to sensory precision, prediction errors, and affective reactivity, pointing to more complex and indirect relationships with pain (99). In (79), gamma was also positively related to alpha activity, which as discussed above is often associated with reduced pain. Broadly, the fact that we observed analgesia with gamma stimulation is inconsistent with a simple, positive relationship between gamma and pain. However, it should also be noted that gamma activity is less easily driven by noninvasive stimulation and may require a specific resonance condition or circuit alignment to influence behavior, which could also explain its weaker effects.

Together, these results support the view that tTIS interacts with intrinsic oscillatory networks and that stimulation is most effective when the applied rhythm matches or amplifies endogenous frequencies that naturally participate in pain regulation. Future studies using EEG or MEG alongside tTIS are needed to gain further insight into the potential mechanisms.

Pain relief was similar on both forearms despite unilateral stimulation of the left M1, with no significant differences between contralateral and ipsilateral sites. This bilateral pattern suggests that M1-targeted tTIS engages body-wide inter-effector regions rather than acting solely through lateralized motor representations, consistent with the SCAN framework (21–25,28). In this framework, inter-effector regions are strongly interconnected both within and across hemispheres and exhibit robust connectivity with control-related regions such as the cingulo-opercular network, supporting coordinated, whole-body action rather than isolated effector-specific control (25). This is notable given that classical corticospinal motor outputs from M1 are strongly lateralized, primarily projecting contralaterally to control limb movements (100), suggesting that the present bilateral effects are unlikely to arise solely from direct motor pathways. Instead, converging evidence from nonhuman and rodent studies indicates that M1 projects broadly to subcortical and bilateral targets, including thalamus, brainstem, and striatal regions, enabling more distributed and bilateral modulation of behavior (21,22). In addition, primate studies show that regions within M1 receive nociceptive spinothalamic input (23) and are involved in coordinating multi-effector motor behavior and autonomic responses, exerting body-wide influence through connections with internal organs and brainstem systems (24,101). Through these connections, stimulation may influence descending pain-regulatory pathways, including projections to the periaqueductal gray and thalamus (22), as well as higher-order networks shaping the cognitive and affective dimensions of pain (see Figure 1).

While left M1 tTIS in this study produced bilateral analgesia that was equivalent on ipsilateral and contralateral arms overall, we did observe some lateralization in the effects across stimulation frequencies. The analgesia difference for 10 Hz vs. higher frequencies was stronger on the ipsilateral forearm, broadly inconsistent with a contralateral motor effect, unless an intra-hemispheric inhibition mechanism is involved (102,103). Several pieces of evidence argue against this interpretation. First, intra-hemispheric inhibition should not produce equivalent overall effects on both sides of the body, but rather predicts opposing effects. Second, the significant interaction between body side and frequency should be interpreted cautiously given the absence of an overall side difference and a non-significant Active vs. Sham × Body Side interaction. Rather than indicating lateralized analgesia, this pattern likely reflects a frequency-specific asymmetry superimposed on an otherwise bilateral effect. Consistent with this interpretation, stimulation did not affect reaction time in the contralateral reporting hand, arguing against a motor or corticospinal mechanism. (103)

The inter-effector concept may also help explain variability in prior M1 stimulation studies. Results from tDCS and tACS targeting M1 have been mixed (18,38), with some studies reporting analgesic effects and others failing to observe reliable changes in pain perception. These inconsistencies are likely driven primarily by methodological factors, including diffuse current spread, limited spatial precision, and reduced ability to engage deeper or distributed cortical–subcortical circuits. While inter-effector regions within M1 have been proposed as potential contributors to body-wide pain modulation, inconsistent findings across studies do not necessarily depend on their selective engagement. Rather, variability may arise more generally from differences in how effectively stimulation approaches modulate relevant motor–cognitive and descending pain-regulatory networks. In this context, tTIS offers several potential advantages, including improved focality, greater effective depth, and frequency-specific modulation of neural activity. These properties may enable more reliable engagement of pain-related networks, providing a mechanistic explanation for the robust and consistent analgesic effects observed in the present study.

Several limitations should be noted. This study focused on acute experimental pain in healthy adults, and further work is needed to determine whether similar effects occur in chronic pain patient populations. Though our results show clear behavioral effects, the underlying neural mechanisms remain to be identified in future studies that combine tTIS with fMRI and EEG or MEG (104,105). A final consideration is that the observed frequency dependent pattern may reflect individual variability in resonance and anatomical features that shape the induced electric field. Future studies that integrate computational modeling, neuroimaging, and individualized stimulation strategies may clarify how these factors influence treatment response (48,49).

Despite these considerations, the present study provides strong initial evidence that tTIS can modulate human pain perception in a frequency-dependent manner. These findings point to a potential fundamental advance in understanding the neural mechanisms underlying human affect and motivation, highlighting how frequency-specific modulation of distributed circuits can shape subjective experience. By altering the dynamics of oscillatory networks involved in sensory and affective processing, tTIS provides a novel tool to causally probe these systems. Looking forward, this approach may enable future applications aimed at understanding and mitigating dysregulation across a range of clinical conditions, including chronic pain and affective disorders, by targeting circuit-level dysfunction with greater precision.

## Conclusion

This study demonstrates that temporal interference stimulation of the primary motor cortex can reliably reduce acute thermal pain in humans, and that the magnitude of analgesia depends on stimulation frequency. All active envelope frequencies reduced pain relative to sham, with the largest effect at 10 Hz, intermediate effects at 20 Hz, and a smaller yet reliable effect at 70 Hz. Pain relief was bilateral and did not decay across the full post-stimulation testing period, and the effects were independent of sensory side effects or expectancy. This bilateral pattern is consistent with the hypothesis that M1-targeted tTIS engaged body-wide inter-effector regions and their downstream pain-regulatory circuits. Together, these findings support a model in which tTIS interacts with intrinsic oscillatory dynamics in sensory and modulatory circuits, with maximal efficacy when stimulation aligns with functionally relevant rhythms. More broadly, frequency-tuned tTIS offers a promising path toward precise, scalable noninvasive neuromodulation for pain, motivating future mechanistic and clinical studies that integrate neuroimaging and individualized field-informed targeting.

## Acknowledgments

We thank Melanie Steiner (TI Solutions AG) for assistance in generating the computational modeling figure illustrating the effects of tTIS on the motor cortex. **Funding:** This work was supported by both the National Institute of Mental Health (No. R37 MH076136; Tor Wager), and the Neukom Institute for Computational Science at Dartmouth College. **Author contributions:** Conceptualization: AD, DMG, TDW; Methodology: AD, DMG, TDW; Investigation: AD, TDW; Visualization: AD; Funding acquisition: AD, EKM, TDW; Project administration: AD, DMG, TDW; Supervision: RJH, TDW; Writing – original draft: AD, DMG, TDW; Writing – review & editing: AD, DMG, EKM, TDW. **Competing interests:** The authors report no conflicts of interest. **Data and materials availability:** All data used for analyses in this study are publicly available in the CANLAB repository at https://github.com/canlab/tTIS_pain_2025.

## Materials and Methods

### Participants

42 healthy participants were recruited from Dartmouth College and the surrounding New Hampshire community. Exclusion criteria were age <18 or >55 years, chronic pain, current or history of major psychiatric or neurological diagnosis, substance abuse in the last 6 months, or having contraindications for brain stimulation or experimental pain (e.g., pregnancy). All participants were also naïve to tTIS, helping to reduce expectancy effects and enhance blinding integrity. The study was approved by the Dartmouth College Institutional Review Board, and informed consent was obtained from all participants before every session, in accord with the Declaration of Helsinki.

Based on findings from a prior study (106), which reported a large effect size (Cohen’s *d* = 0.86) for the contrast between anodal and cathodal tDCS in pain modulation, we estimated a medium effect size (*d* = 0.6) or larger for the contrast between active and sham stimulation in the current design. Assuming a one-tailed test at α = 0.025, a sample size of N = 32 was required to detect this effect with 90% power. We initially recruited 42 participants. Six participants were excluded based on their pain calibration results, and four participants dropped out of the study during or after the first tTIS session, leaving a final sample size of N = 32 (16 females, M_age_ = 24.7 ± 4.6, range: 18–34 years). The study design is presented in Figure 2.

### Adaptive Thermal Calibration Procedure (Session 1)

Participants first completed an adaptive thermal calibration procedure to individualize stimulus intensity and estimate pain threshold and tolerance (full methodological details in (32,106). Calibration was conducted on the volar forearm using six predefined stimulation sites based on anatomical landmarks (Figure 2b).

The calibration consisted of four runs (24 trials total). Run 1 served as a familiarization phase and was excluded from analysis. After each heat stimulus, participants rated sensation intensity using a gLMS (107) and then provided two separate “yes/no” responses indicating whether the stimulus was painful and tolerable. The gLMS was designed to have ratio scaling properties, facilitate cross-sensory comparisons, and avoid ceiling effects with intense sensations, and included anchors ranging from “No sensation” (0°) to “Barely detectable” (3° of scale angle), “Weak” (10°), “Moderate” (29°), “Strong” (64°), “Very Strong” (98°), and “Strongest sensation of any kind” (180°). Run 2 consisted of fixed-temperature stimuli, with 45°C, 47°C, and 48.9°C each applied twice in random order, to initialize logistic regression models estimating pain threshold and tolerance. Runs 3 and 4 implemented an adaptive procedure in which stimulus temperatures from the logistic regression models were updated on each trial and randomly assigned to low and high categories (three trials each per run), selecting temperatures associated with a fitted probability of 0.9 for exceeding the pain threshold or remaining below the tolerance threshold, thereby sampling near individual pain and tolerance boundaries and allowing progressive refinement of threshold estimates.

Following calibration, a linear regression model was used to estimate pain intensity as a function of temperature. The two skin sites with the lowest mean absolute prediction error were selected for subsequent sessions, ensuring comparable pain sensitivity across stimulation sites. Participants whose estimated pain threshold and tolerance differed by less than 1°C were excluded. 42 participants completed the calibration session and 36 participants passed and were selected for inclusion in the tTIS sessions.

### Experimental Sessions (Sessions 2-4)

Participants returned for four additional sessions on separate days, with a minimum three day interval to minimize carryover effects of tTIS (8.5 ± 6.4, range: 3–43 days between sessions), consistent with prior studies using multi-day washout periods in crossover designs (108–110). In each session, participants underwent four eight-minute pain tests at two different skin sites on the right and left forearm (pre-tTIS, two runs per forearm side, 12 trials per skin site, 48 trials total), followed by 20 minutes of continuous tTIS, a questionnaire about the tTIS experience, and finally four eight-minute pain runs at two different skin sites on the right and left forearm (post-tTIS, two runs per forearm side, 12 trials per skin site, 48 trials total). The four pre- and post-tTIS runs were conducted back-to-back with no planned interval between them; the only pause was the ∼30–60 seconds required to move and secure the thermode to a new skin site. The order of stimulation across left and right forearms, as well as the sequence of skin sites, was randomized across runs. tTIS was either sham, 10, 20, 70 Hz over the left M1, with participants randomly assigned to a treatment order (Figure 2a).

### Noxious Heat Stimulation

Noxious heat was delivered randomly to two locations on the right and left forearm using a TSA-II Neurosensory Analyzer (Medoc Ltd) with a 16-mm thermode end-plate. Participants received four 8-min runs of repeated stimulation before and after tTIS stimulation (eight runs total). Each run included 12 trials with 6 randomly individually calibrated low and 6 high temperatures (see Results for intensities) delivered to the right and left volar forearm. Each trial lasted for 40 seconds and started with a warning cue (the words “Get Ready”; 1 second), followed by a 2-6 second delay, heat stimulation (13 seconds including 1.5 seconds ramp up/down, 10 seconds plateau), a 5-9 second delay, a pain rating period (7 seconds using right hand), and a final 4-12 second delay before the next cue. Participants rated the sensation using the semi-circular gLMS with their left hand.

### TTIS Design

TTIS was administered using the Temporal Interference Brain Stimulator for Research (TIBS-R v3.0, TI Solutions AG, Switzerland). Electrode positioning was guided by a computational modeling platform developed by the device manufacturer, which optimized current delivery to selectively target the left motor cortex. The optimized electrode montage is shown in Figure 2(c) and the electrode pairs are F3-P3 and FC1-CP1.

Stimulation was delivered via two independent electrode pairs, with each circular Ag/AgCl electrode having a surface area of 3.14 cm². Electrodes were embedded in an EEG cap provided by Neuroelectrics and affixed to the scalp using EEG gel to ensure low-impedance contact.

Each electrode pair delivered 2 mA of sinusoidal current, for a total of 4 mA across both channels. Temporal interference was generated using two high-frequency carrier signals (1 KHz) (48) differing by 10 Hz, 20 Hz, or 70 Hz, corresponding to the desired beat frequency. These beat frequencies were selected to engage neural oscillations within alpha, beta, and gamma bands, respectively.

Stimulation was applied for 20 minutes, with a 10-second ramp-up and ramp-down period to minimize perceptual sensations. Electrode impedance was checked prior to and during the experiment, and maintained at below 5 kΩ to ensure consistent contact quality.

A sham stimulation condition was included as a control. During sham, stimulation ramped up over 10 seconds, was held at peak amplitude for 1 minute, and ramped down over 10 seconds. The beat frequency during sham was randomly selected from 10, 20, or 70 Hz to maintain sensory similarity while avoiding sustained neuromodulatory effects.

### Analysis Model

The study included a 4 x 2 experimental design (tTIS Type x Stimulus Intensity: Sham, 10, 20, and 70 Hz x low and high temperature from calibrated pain and tolerance thresholds). We tested the main effects of each factor and their interaction, with planned contrasts capturing differences across tTIS conditions (Sham > 10 & 20 & 70, 10 > 20 & 70, and 20 > 70 Hz), as well as the durability of tTIS effects over time, potential effects of stimulation order, tTIS induced sensations (e.g., itching, tingling), and other effects described below. Thermal sensation ratings that deviated more than 3 standard deviations from a participant’s mean value within each of the 8 cells in the 4 x 2 design were excluded. Less than 1% of trials were excluded based on this criterion.

The statistical model was a mixed-effects model with random participant intercepts and random slopes for tTIS Type, Stimulus Intensity, and tTIS Type x Stimulus Intensity (low and high temperature). This model was implemented using the “fitlme” function in MATLAB R2024b (MathWorks, Natick, MA, USA) with maximum likelihood parameter estimation. For this purpose, full trial-level data were included as well as covariates including pre-treatment mean pain scores and treatment order groups (4 groups; see Figure 2a). This mixed-effects model tests main effects and interactions analogous to repeated-measures ANOVA, but is preferred to ANOVA on participant-level summary statistics because it models the within-subject covariance structure directly rather than assuming sphericity. This typically improves efficiency (power) and yields more valid inferences when sphericity is violated or variance–covariance structure is heterogeneous (111,112). We considered additional covariates for sensations elicited by tTIS Type (e.g, itching, tingling, metallic-iron taste, pain, warmth, and pleasantness), but sham, 10, 20, and 70 tTIS produced equivalent self-reported sensations (all p > 0.05), so we did not include these covariates in the main model. The Satterthwaite method was employed to estimate the degrees of freedom. In addition, to screen for baseline biological sex differences, we compared calibrated pain and tolerance thresholds between biological females and males using independent-samples Welch’s t-tests. The mixed-effects model used in this study is summarized as:

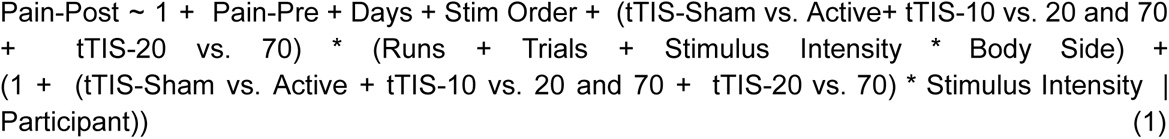

In Equation 1, Pain-Post is the pain rating on each trial after tTIS treatment, tTIS-Sham vs. Active represents the effect of sham vs. average effect of 10, 20, and 70 Hz stimulation (coded with [3/4, -1/4, -1/4, -1/4] for sham, 10, 20, and 70 Hz), tTIS-10 vs. 20 and 70 represents 10 vs. average effect of 20 and 70 Hz (coded with [0 2/3 -1/3 -1/3] for sham, 10, 20, and 70 Hz), and tTIS-20 vs. 70 represents the effect of 20 vs. 70 Hz stimulation (coded with [0, 0, 1/2, -1/2] for sham, 10, 20, and 70 Hz). Pain-Pre is the mean pain rating before tTIS in each session. Stim Order refers to four different stimulation orders across four tTIS sessions (see Figure 2 for four different orders across tTIS sessions). Days, Runs, and Trials refer to linear effects of session (testing day, 4 in total, mean-centered), run number (run within session across 4 runs, mean-centered within day), and trial number (trial within run across 12 trials, mean-centered within run), respectively. These variables were included to control for habituation and/or sensitization across time. Body Side indicates right vs. left forearm stimulation, Stimulus Intensity represents the temperature obtained by calibration (low and high temperature based on pain and tolerance threshold, coded with 1 and -1 for high and low), and Participant represents the random effect of the participant. In this model the interaction between tTIS Type and Runs and tTIS Type and Trials within run test the effect durability of tTIS over the 40-min post-stimulation test period. To evaluate the robustness of the primary findings, we conducted a series of supplementary mixed-effects analyses. These analyses included models that additionally adjusted for age and biological sex, expanded the random-effects structure by incorporating session effects and allowing trial- and run-level slopes to vary by tTIS condition, and controlled for self-reported tTIS-induced sensations (e.g., itching, tingling). We also re-estimated the primary model after excluding the first pain trial to account for potential novelty or orienting effects at the start of each run.

## Supplementary materials

The complex model used in the first sensitivity analysis is presented in this section.

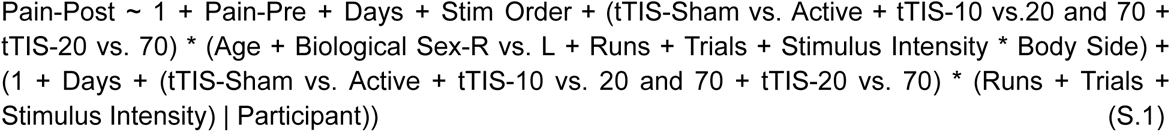

Pain rating changes for high-temperature (Figure S1a) and low-temperature (Figure S1b) conditions across all stimulation types are presented here.

**Figure S1.**
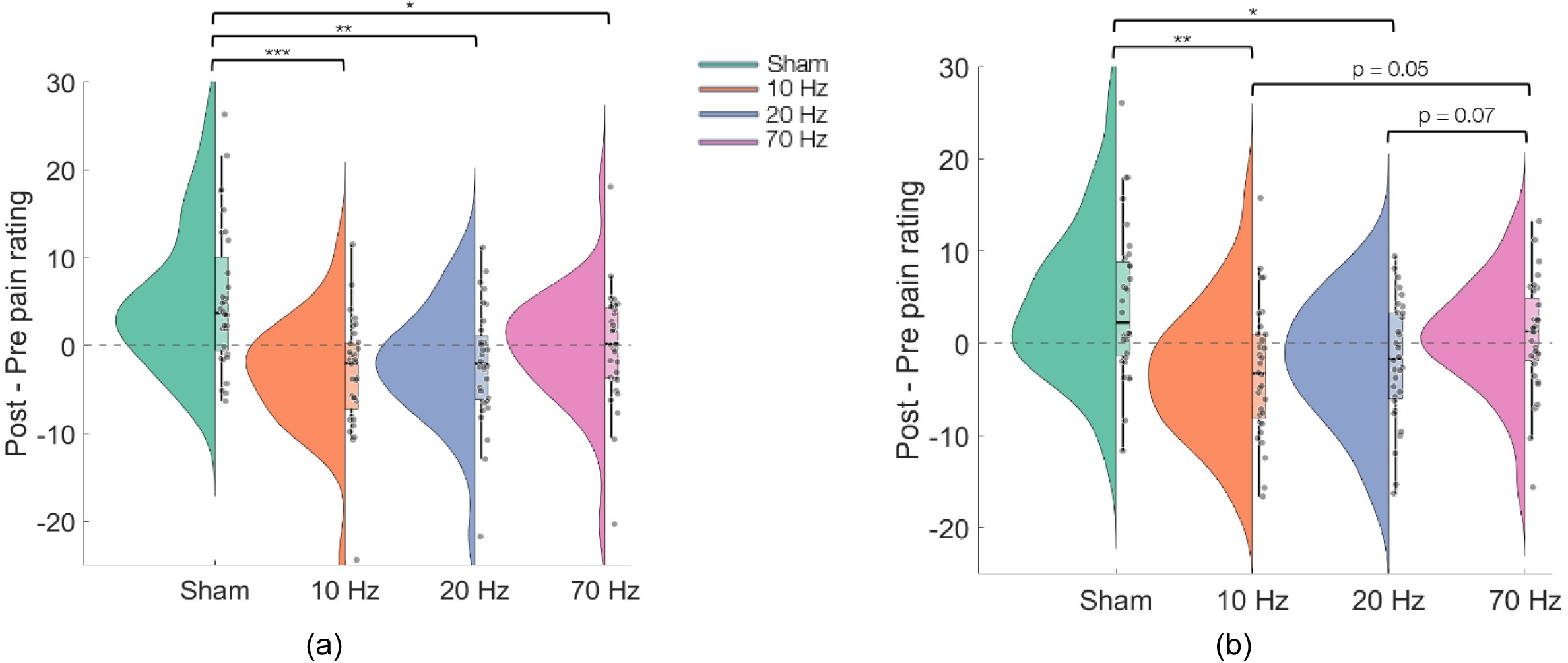
Main findings for pain sensation. (a-b) Treatment effects (Post-treatment—Pretreatment pain rating change) for each of the 4 stimulation types and for each of high temperature (left panel) and low temperature (right panel). Boxplots indicate the median, 25th, and 75th percentiles, and individual data points. Dots reflect effect magnitude estimates of individual participants. All values were adjusted to remove participant-level intercepts. \*\*\**P* < 0.001; \*\**P* < 0.01; \**P* < 0.05 two-tailed.

## Notes

### Competing Interest Statement

The authors have declared no competing interest.

