## Supplemental Materials for "Temporal Interference Stimulation of the Motor Cortex Produces Frequency-Dependent Analgesia"

**Supplementary materials**

The complex model used in the first sensitivity analysis is presented in this section.

Pain-Post ~ 1 + Pain-Pre + Days + Stim Order + (tTIS-Sham vs. Active + tTIS-10 vs.20 and 70 + tTIS-20 vs. 70) * (Age + Biological Sex-R vs. L + Runs + Trials + Stimulus Intensity * Body Side) + (1 + Days + (tTIS-Sham vs. Active + tTIS-10 vs. 20 and 70 + tTIS-20 vs. 70) * (Runs + Trials + Stimulus Intensity) | Participant)) (S.1)

Pain rating changes for high-temperature (Figure S1a) and low-temperature (Figure S1b) conditions across all stimulation types are presented here.

**Figure S1. Main findings for pain sensation.**
(a-b) Treatment effects (Post-treatment—Pretreatment pain rating change) for each of the 4 stimulation types and for each of high temperature (left panel) and low temperature (right panel). Boxplots indicate the median, 25th, and 75th percentiles, and individual data points. Dots reflect effect magnitude estimates of individual participants. All values were adjusted to remove participant-level intercepts. ***$P < 0.001$; **$P < 0.01$; *$P < 0.05$ two-tailed.

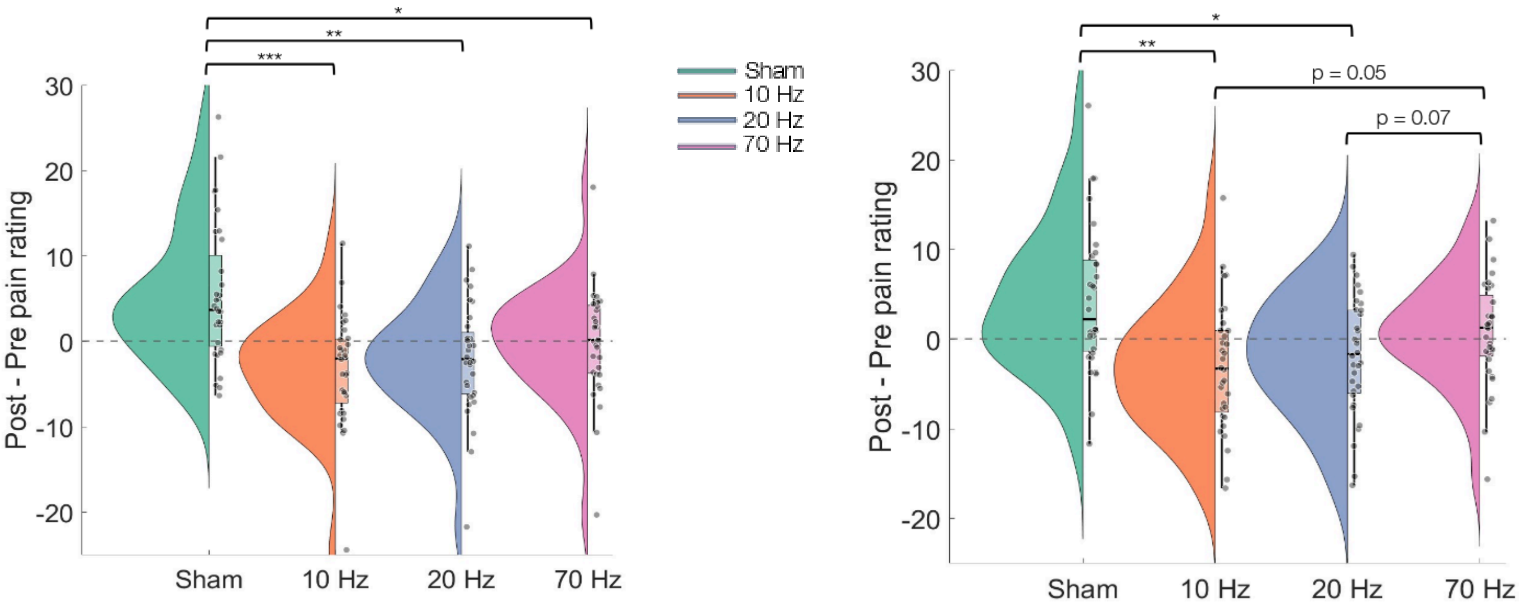


(a) (b)
